# Connectional heterogeneity in the mouse auditory corticocollicular system

**DOI:** 10.1101/571711

**Authors:** Georgiy Yudintsev, Alexander Asilador, Macey Coppinger, Kavyakrishna Nair, Masumi Prasad, Daniel A. Llano

**Author notes:** Current address: Department of Microbiology, University of Georgia, Athens, GA, 30602.

## Abstract

The auditory cortex (AC) sends long-range projections to virtually all subcortical structures important for hearing. One of the largest and most complex of these - the projection between AC and inferior colliculus (IC, the corticocollicular pathway) - has attracted attention due to its potential to alter IC response properties. The corticocollicular pathway comprises a component originating from layer 5, but recent evidence suggests a significant contribution from deep layer 6, constituting 25% of corticocollicular neurons in mouse. The functions of layer-specific corticocollicular projections are poorly understood. Here, using a combination of tracers and *in vivo* imaging, we observed that layer 5 and layer 6 corticocollicular neurons differ in their cortical areas of origin, as well as IC termination patterns. Layer 5 corticocollicular neurons are concentrated in primary AC areas while layer 6 corticocollicular neurons emanate from broad auditory and non-auditory areas of temporal cortex. In addition, layer 5 projects to three IC subdivisions with axo-somatic terminals in the central nucleus, while layer 6 projects to non-central nuclei and targets the most superficial layers. These findings suggest that layer 5 corticocollicular neurons form a direct connection between primary AC and IC while the layer 6 projection is more diffusely organized and carries non-auditory information to modulate IC.

## Introduction

The central auditory system consists of a series of brainstem, midbrain and thalamic nuclei, and neocortical regions. The classical ascending central auditory pathway begins with the cochlear nucleus (CN), from which the information about sound reaches the superior olivary complex (SOC), followed by the nuclei of the lateral lemniscus (NLL), the inferior colliculus (IC), the medial geniculate body of the thalamus (MGB), and, finally, the auditory cortex (AC) (Malmierca and Ryugo 2012). A prominent feature of the central auditory system is the presence of massive descending connections, which arise virtually at all levels of the auditory pathway (Winer 2005, Bajo, Nodal et al. 2007, Bajo, Nodal et al. 2010, Terreros and Delano 2015, Lesicko and Llano 2017, Patel, Sons et al. 2017). One of these descending pathways, the pathway between the AC and IC, referred to here as the corticocollicular system, has recently garnered much attention due to its potential to alter the sensory information processing at the level of the IC. For example, electrical stimulation of frequency-specific regions of the AC was shown to evoke immediate and long-lasting changes of best frequency representation in the IC in a corticocentric fashion (Yan and Suga 1998, Yan, Zhang et al. 2005). In addition to altering of the frequency representation, the neurons in the IC also exhibited changes in tuning to a number of other sound features including duration, intensity and location (Ma and Suga 2001, Yan and Ehret 2002, Zhou and Jen 2005). The corticocollicular system has also been implicated in mediating experience-induced auditory plasticity (Bajo, Nodal et al. 2010), control of an innate sound-evoked escape behavior (Xiong, Liang et al. 2015), and compensatory gain changes following a significant loss of peripheral auditory input (Asokan, Williamson et al. 2018). Such functional heterogeneity is unlikely to be supported by a single type of the corticocollicular neuron, and calls for further understanding and characterization of the anatomy of this projection system. Despite a mounting body of research elucidating the functions of the corticocollicular system, a detailed picture of the functional neuroanatomical organization of this pathway remains to be uncovered.

The auditory corticocollicular projections emanate from distinct regions of layer 5 and lower layer 6 of the temporal cortex. Dual layer 5/layer 6 projections have been documented across multiple species (Games and Winer 1988, Künzle 1995, Coomes, Schofield et al. 2005, Bajo, Nodal et al. 2007, Schofield 2009), and neurons from these two layers possess different morphological, electrophysiological and local circuit properties (Slater, Willis et al. 2013, Zurita, Rock et al. 2018, Slater, Sons et al. 2019). In a related descending system, the corticothalamic system, layer 5 and layer 6 projections also have distinct morphological, electrophysiological and network properties (Llano and Sherman 2008, Llano and Sherman 2009), and have been hypothesized to have different roles in modulating the thalamus and supporting cortico-cortical communication (Ojima 1994, Guillery and Sherman 2002, Theyel, Llano et al. 2010). Whether a similar functional distinction exists between layer 5 and layer 6 corticocollicular projections is not yet known.

One key set of questions regarding layer 5 and layer 6 corticocollicular projections is whether they originate from different regions of the cortex or terminate differentially in the IC. Early neuroanatomical experiments described regional distributions of layer 5 corticocollicular neurons with respect to the IC, but used insensitive retrograde tracers that did not label the layer 6 pathway (Herbert, Aschoff et al. 1991).

In the present study, using sensitive tracers and molecular tools to separate layer 5 from layer 6 projections as well as in vivo imaging to identify cortical regions, the distributions and cortical regions of origin of layer 5 and layer 6 corticocollicular neurons in the mouse were examined. Substantial heterogeneity and a regional non-overlap between the corticocollicular neurons arising from the two layers was found. Layer 5 corticocollicular neurons were concentrated over a smaller area of the mouse AC, largely confined to its primary regions, while the areal distribution of layer 6 corticocollicular neurons was wider, extending beyond non-primary mouse AC regions and into non-auditory regions. Such neuroanatomical organization may partially explain the functional heterogeneity observed in *in vivo* studies of the corticocollicular system, and provides a foundation for forming and testing future hypotheses about this descending projection system.

## Materials and Methods

### Animal preparations and surgical procedures

All surgical procedures were approved by the Institutional Animal Care and Use Committee at the University of Illinois at Urbana-Champaign. Mice were housed in animal care facilities approved by the American Association for Accreditation of Laboratory Animal Care (AAALAC). Every attempt was made to minimize the number of animals used and to reduce suffering at all stages of the experiments. For anatomical experiments and reconstructions, adult (60-90 days) and young adult (45-60 days) BALB/c mice of both sexes were used. Animals were anesthetized with ketamine hydrochloride (100 mg/kg) and xylazine (3 mg/kg) intraperitoneally and placed in a stereotaxic apparatus (David Kopf Instruments, Tujunga, CA). Aseptic conditions were maintained throughout the surgery. Fluorogold (Fluorochrome, LLC, Denver, CO) was either pressure-injected, or injected into the IC via iontophoresis. For pressure injections, Fluorogold was dissolved (1%) in distilled water and 500 nL was pressure-injected into the left IC using glass pipettes with 10-14 μm in diameter. For iontophoretic injections, Fluorogold was dissolved in acetate buffer (0.1 M) at pH 3.4, and injected into the left IC using iontophoresis through a 20 μm tip diameter broken glass electrode for 10-15 minutes at 10 μA positive current, with 7 s on 7 s off (50% duty cycle). These protocols resulted in large unilateral injections into the left IC. For in vivo imaging experiments (see details below) and reconstructions, C57BL6J-Tg(Thy1-GCaMP6s)GP4.3DkimJ mice were purchased from Jackson Laboratories, bred as heterozygous in the local animal facility. Prior to the experiments, mice were genotyped for the presence of Tg(Thy1-GCaMP) transgene (forward PCR primer CAT CAG TGC AGC AGA GCT TC, reverse PCR primer CAG CGT ATC CAC ATA GCG TA). Before the start of the surgery to prepare mice for imaging, mice were anesthetized with a mixture of ketamine and xylazine (100 mg/kg and 3 mg/kg respectively) delivered intraperitoneally using a 27-gauge needle, following by an intraperitoneal injection of acepromazine (2-3 mg/kg). A skin incision over the dorsal portion of the skull was made, after which the skull was exposed. The dorsal surface of the skull was roughened using a small surgical drill bit. The area was cleaned of any remaining bone pellets, and a small aluminum bolt of 1 cm in length was secured to the surface using dental cement (3M ESPE KETAC). When the cement is set, the animal was carried into a dark sound-proof chamber for imaging. Proper care was taken to maintain body temperature within the range of 35.5 to 37°C during imaging using a DC temperature controller (FHC, ME, USA) and a rectal thermometer probe.

Upon the completion of each imaging session, Fluorogold was injected into the left IC of these mice, as above. Three fiducial markers were created by iontophoresis of small amounts of tetramethylrhodamine 10,000 MW dissolved in PBS (Invitrogen, Grand Island, NY) into the cortical regions outside the AC. The iontophoretic injections were done using unbroken glass electrode (tip size ∼ 0.5 μm) with 5-μA positive current and 7 s 50% duty cycle, for 4 minutes. Using the same settings as during *in vivo* imaging, a micrograph of the skull surface and vasculature was taken to aid in co-registration of functional maps with the reconstructions in Neurolucida later. Following the imaging experiment and the surgery, animals were allowed to survive for 7 days, after which the brain tissue was processed according to a standard histological protocol described further below.

For AAV injections, the left ACs of Rbp4-positive mice (n = 2) were injected with 250-500 nL of a mixture of equal concentrations of AAV9.CB7.CI.mCherry.WPRE.rBG (a Cre-independent virus) and AAV9.CAG.Flex.eGFP.WPRE.bGH (a virus only expressed in Cre+ cells), both at 3.43×10^12^vg/mL, Penn Vector Core, Philadelphia, PA). In one Rbp4+CLGT+ mouse, a Cre-OFF virus (pAAV-Ef1a-DO-hChR2(H134R)-mCherry-WPRE-pA) was injected to label layer 6 corticocollicular terminals. Same surgical protocol and post-surgical procedures were applied as described above.

For retrograde tracer injections into the cortex, 250 nL of Alexa-594 conjugated cholera toxin subunit b (recombinant), CTb (0.5 % in distilled water) (Thermo Fisher Scientific, Grand Island, NY) was pressure-injected according to the protocol described above.

### Macroscope and in vivo imaging set-up

An Imager 3001 Integrated data acquisition and analysis system (Optical Imaging Ltd., Israel) was used to image the cortical responses to sound in C57BL6J-Tg(Thy1-GCaMP6s)GP4.3DkimJ mice. A macroscope consisting of 85 mm f/1.4 and 50 mm f/1.2 Nikon lenses was mounted to an Adimec 1000m high-end CCD camera (7.4 × 7.4 μm pixel size, 1004 × 1004 pixels, 10 frames/s), and centered above the left AC, focused approximately 0.5 mm below the surface of the exposed skull. Blue excitation (450 nm, 30 nm band-pass), green emission (515 nm, long-pass) filters and a 495 DRLP dichroic mirror were used. Imager 3001 VDAQ software controlled the acquisition and stimulus trigger.

### Acoustic stimulation

Acoustic stimuli were generated using a TDT System 3 with an RP 2.1 Enhanced Real-Time Processor and delivered via an ES1 free field electrostatic speaker (Tucker-Davis Technologies, FL, USA), located 8 cm away from the contralateral ear. All imaging experiments were conducted in a soundproof chamber. 500 ms pure tones of 5, 10, 20, and 30 kHz were used, 100% amplitude-modulated at 20 Hz. In another set of experiments, a series of species-specific mouse calls were used as auditory stimuli. The recordings were used by previous investigators (Grimsley, Monaghan et al. 2011, Grimsley, Sheth et al. 2016), and kindly made available by this group. In this study, four calls from three major groups were used. Two calls (call 1 and 2) are from a group of low-frequency stress calls. Another call (call 3) is a medium-frequency stress call, which the animals produce when restrained. The final call used (call 4), is a mating call. All playbacks were sampled at 200 kHz. Call 1, 3 and 4 are 500 ms in duration, call 2 is 300 ms.

### Analysis of in vivo imaging activity

Custom-written MATLAB software was used to obtain *ΔF/F*-responses to pure tones (5, 10, 20, and 30 kHz) and species-specific mouse calls. In this way, multiple AC regions could be identified. 2.5 standard deviations threshold was used as a cutoff point to display the peaks of neural signals signal.

### Fluorescence microscopy and reconstructions

All coronal sections of the brain that contained retrograde label were serially photographed using an Olympus IX71 epifluorescence microscope using 5x 0.15 NA objective. These images were then used to create 3-D reconstructions of the left AC, which was done in Neurolucida (MBF Bioscience, Williston, VT). For counting cells, a grid consisting of twelve rectangular bins (bin size 118,360 μm^2^) was placed throughout images of the cortex in each coronal section, such that all layers of the cortex were covered. The rhinal fissure was used as a reference point for grid placement, with two bins of the grid being positioned ventral to the rhinal fissure and ten dorsal. Confocal pictures were taken on a Leica SP8 UV/Visible Laser Confocal Microscope. 488 nm and 561 nm excitation was used for visualizing eGFP and mCherry, respectively. For images used for the analysis of terminal size, 500 nm z-step was used.

### Immunohistochemistry

The mice were first deeply anesthetized with a lethal intraperitoneal injection of ketamine hydrochloride (200 mg/kg) and xylazine (6 mg/kg), and perfused transcardially with 4% paraformaldehyde in phosphate-buffered saline (PBS) at pH 7.4. Frozen 50 µm sections were cut using a sliding microtome. Prior to cutting, three fiducial markers were placed in brain tissue along the rostrocaudal axis using a 27-gauge needle dipped into water-insoluble black India ink. The fiducials were placed to ensure alignment of serial coronal sections for reconstructions in Neurolucida. For immunohistochemistry, sections were washed 3 times for 5 minutes in PBS, microwaved for 15 seconds at full power for antigen retrieval, then incubated for 30 minutes in PBS containing 0.3% Triton-X (PBT) followed by a 30 minute blocking step in a 3% serum containing PBT solution. Blocking serum was of the same species that the secondary antibody was generated in. Sections were then incubated with corresponding primary antibody overnight in the cold room. For parvalbumin (Sigma, St. Louis, MO), 1:1000 dilution was used, and for SMI32 (BioLegend, San Diego, CA), the dilution was 1:1500. For NeuN staining, rabbit anti-NeuN antibody (Millipore Sigma, Burlington, MA) at 1:1000 dilution was used. Secondary antibody was diluted in the serum solution used for the primary wash and sections were incubated in this solution for 2 hours at room temperature. To avoid any spectral overlap with Fluorogold during imaging of the sections, Alexa-568 conjugated secondary antibody was used (Invitrogen, Grand Island, NY) to reveal SMI32 or parvalbumin immunoreactivity, and Alexa-633 conjugated secondary antibody for was for NeuN (Invitrogen, Grand Island, NY). Then, the sections were washed in PBS three times for 10 minutes each, mounted on gelatin-coated slides, air-dried in dark room, coverslipped using fluorescence mounting medium (Vectashield H-1000, Vector labs) and sealed with nail polish.

### Analysis of layer 5 and 6 neuronal distributions

Binned cell counts for each layer were exported as Excel spreadsheets. Abercrombie adjustments were done on each layer cell count separately (Abercrombie 1946). Full width at half maximum analysis of neuronal distributions along dorsoventral and rostrocaudal axes was performed. To compute full width at half maximum, cell counts for each layer were summed separately along the rows and along the columns to obtain distributions of counts along the dorsoventral and rostrocaudal axes, respectively. A fourth degree polynomial provided the best fit for each distribution. The peak and full width at half maximum measures were obtained for each layer in both dimensions (dorsoventral and rostrocaudal), using the polynomial fit. This was done for each animal separately. For qualitative comparisons, the individual distributions for layer 5 and 6 were imported into MATLAB (MathWorks, Natick, MA) and plotted as contour maps.

### Statistical analysis

Pairwise differences were analyzed using non-parametric statistical tests. These tests were chosen because of our smaller sample size. Not relying on the assumptions of normality of the underlying sampling distributions, and these tests provide more accurate statistical results. Wilcoxon rank-sum test was used to compare the differences between layer 5 and layer 6 measures of peaks and full-width-half-max measures. Kolmogorov-Smirnov test was used to compare the distributions of the corticocollicular terminals sizes within the IC subdivisions.

## Results

Fluorogold was injected into the left IC of 10 mice, producing retrogradely-labeled corticocollicular neurons in cortical layers 5 and 6 in all animals (Figure 1, A and E). In six cases, the sections were also processed for parvalbumin (PV) or SMI32 fluorescent immunostaining (Figure 1, B and F). Previous studies reported that PV and SMI32 immunoreactivity in the AC delineate lemniscal auditory areas (primary auditory cortex A1 and anterior auditory field AAF (Cruikshank, Killackey et al. 2001, Horie, Tsukano et al. 2015)). However, no studies described the distribution of layer 5 and 6 corticocollicular neurons with respect to these neurohistological markers. Using 50% threshold for fluorescence intensity (Figure 1, C and G), it was observed that the majority of layer 5 corticocollicular neurons were restricted to the A1 and AAF, while layer 6 corticocollicular neurons, particularly more rostrally, appeared in both PV and SMI32 strongly and weakly staining regions, suggesting that certain cerebral cortical regions are connected with the IC only via projection neurons emanating in layer 6 (Figure 1, D, H, and insets K and L).

**Figure 1.**
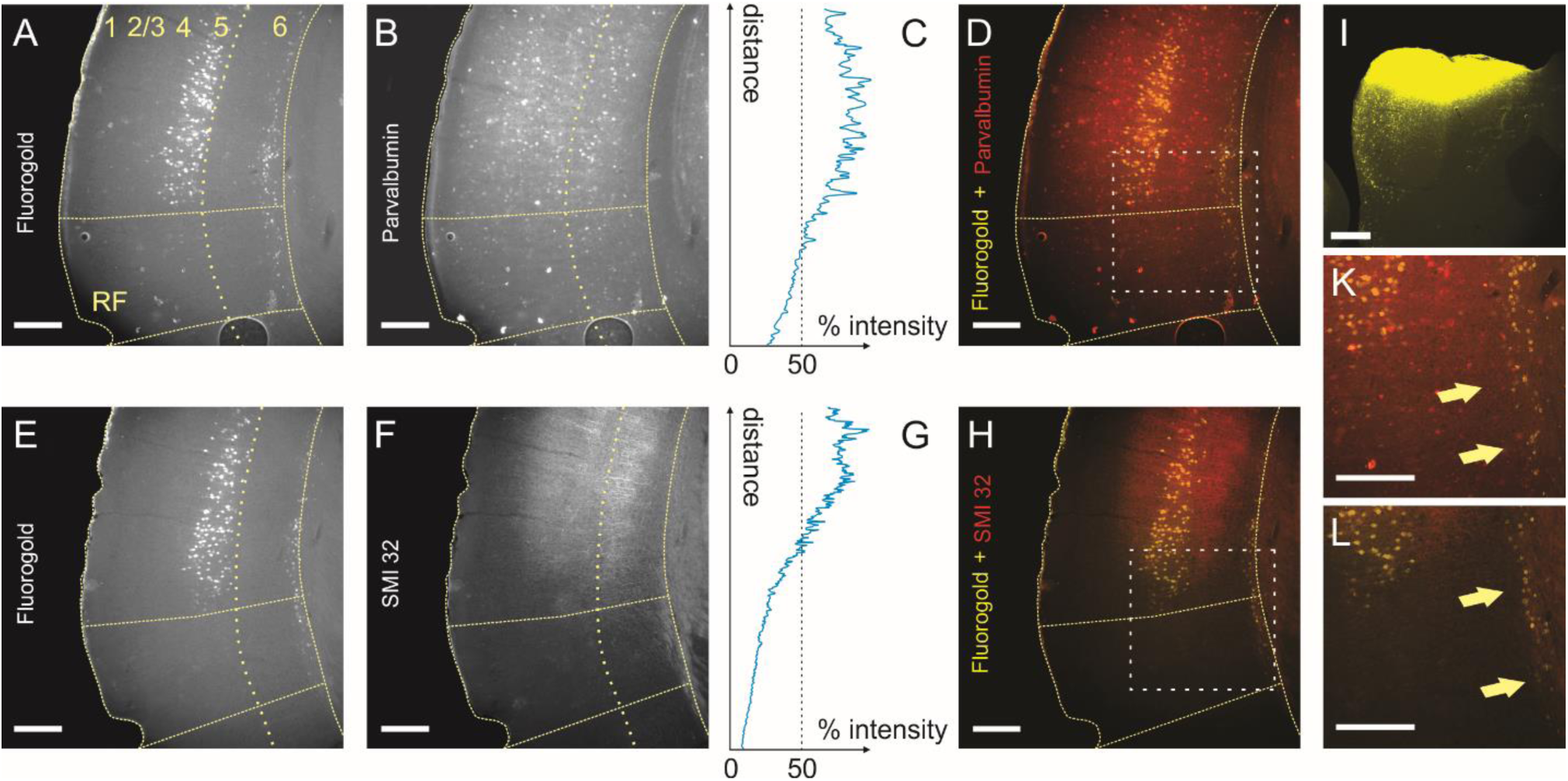
Partially segregated distributions of layer 5 and layer 6 corticocollicular neurons in mouse AC. Corticocollicular neurons in layers 5 and 6 from adjacent coronal sections labeled with Fluorogold (A and E). Parvalbumin (B) and SMI32 (F) immunoreactivity in mouse AC. Fluorescence intensity profiles for parvalbumin (C) and SMI32 (G). Layer 5 corticocollicular neurons are confined to parvalbumin-and SMI32-rich regions of the mouse AC (D and H), while many layer 6 corticocollicular neurons in found in parvalbumin and SMI32 negative cortical regions (D and H, also K and L). (I). Corresponding injection of Fluorogold in the left IC. Scale bar = 250 μm.

To determine whether layer 5 and layer 6 corticocollicular neurons have different distributions, we first plotted all layer 5 and 6 corticocollicular neurons in Neurolucida software so that the backlabeled cells can be visualized in lateral view, similar to Herbert et al. (1991). Micrographs of coronal sections containing corticocollicular neurons were serially aligned and used to create a 3-D reconstruction with plots of layer 5 and 6 corticocollicular cells (Figure 2 A and B). Figure 2 shows the final result of one such reconstruction, where layer 5 corticocollicular cells are represented as red markers, and layer 6 corticocollicular cells as the yellow ones. It appeared that layer 5 and 6 corticocollicular cells were substantially non-overlapping; an area containing layer 6 corticocollicular neurons without any layer 5 corticocollicular neurons was observed rostroventrally.

**Figure 2.**
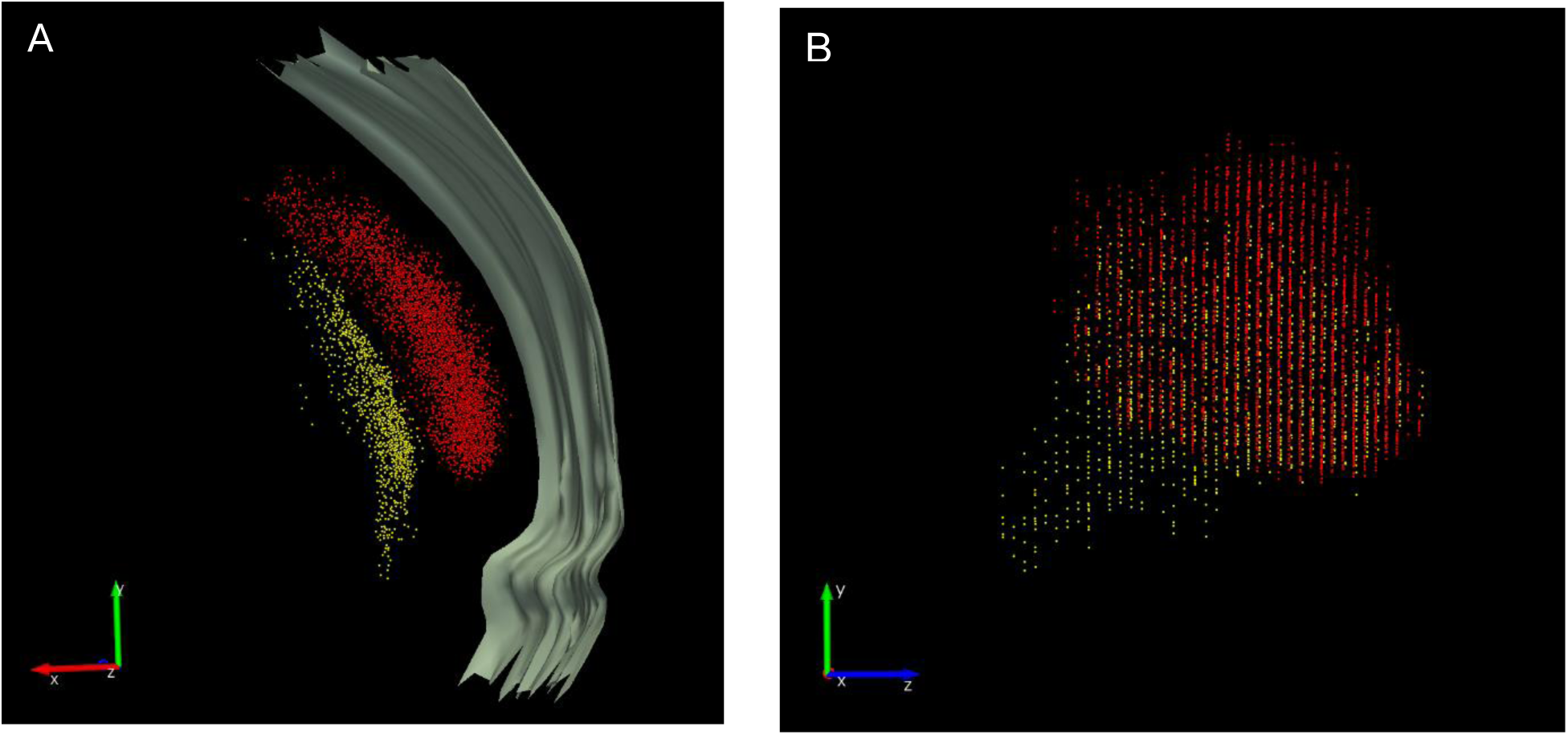
A representative reconstruction of corticocollicular cellular distributions in the cortex. Corticocollicular neurons were marked with a red (layer 5) or a yellow (layer 6) marker. (A) A series of pooled reconstructed coronal sections of the mouse brain on the left side. The outer grey border marks the pial surface. Rotating this reconstruction 90 degrees along the y-axis results in the lateral view of the reconstructed distributions (B). Scale bar = 250 μm.

For further qualitative and quantitative analyses, the distributions data were plotted as contour maps. Figure 3 shows a panel where the rows correspond to individual animals, and the columns contain the injection sites of Fluorogold in left ICs, layer 5 and layer 6 maps, and finally the difference between layers 6 and 5. Here, layer 5 cell counts were subtracted from layer 6, and any only positive values were shown to show regions where the number of layer 6 cells exceeded layer 5. This subtraction revealed cortical areas containing larger numbers of layer 6 corticocollicular neurons in each animal. Qualitatively, it appeared that layer 6 corticocollicular neurons occupy overall a broader area in the cortex compared to layer 5 corticocollicular neurons. In each animal, irrespective of the IC injection site, an area of non-overlap with isolated layer 6 corticocollicular neurons was observed rostroventral to the AC (right-most column in Figure 3). To quantify these differences in distributions, the widths and peaks of the distribution for layers 5 and 6 within each animal (n = 10) were then compared.

**Figure 3.**
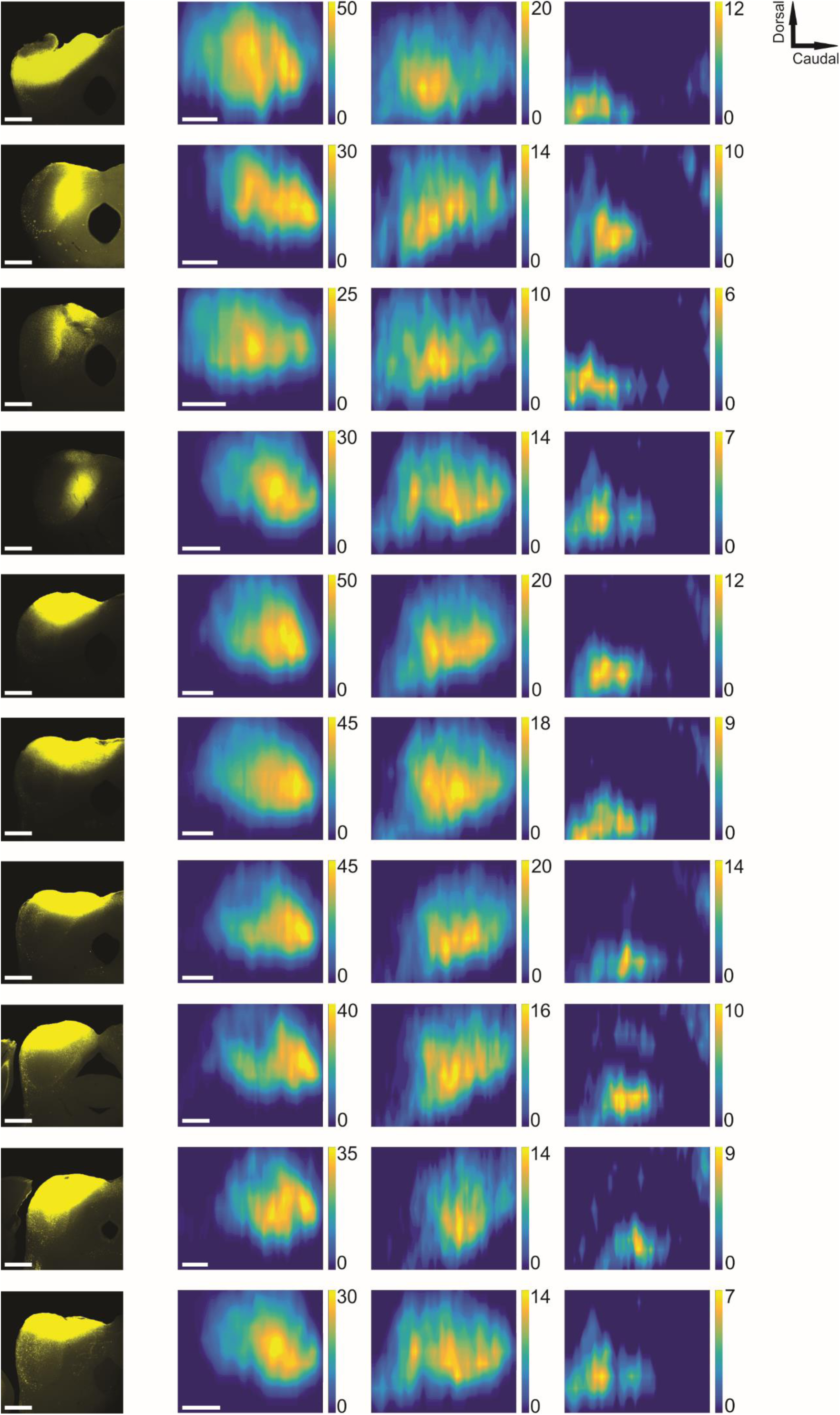
Spatial distributions of layer 5 and 6 corticocollicular neurons. Each row includes data from one animal. The first column includes the injection sites of Fluorogold into the left IC of each animal. The second column shows the distribution of layer 5 corticocollicular neurons in the cortex, while the third column has the corresponding distributions of layer 6 corticocollicular neurons. The final column shows the difference (Layer 6 – Layer 5). The compass in the top right corner describes the directionality for the maps. Scale bar = 250 μm.

The cell counts for each layer and each animal were collapsed in rostrocaudal and dorsoventral axes to obtain two distributions (Figure 4 A and B). Along the dorsoventral axis, the average full-width at half maximum for layer 5 was 824.6 μm (SD = 57,5 μm) and 929.0 μm (SD = 92.1 μm) for layer 6 (n = 10, p-value = 0.0059, Wilcoxon signed rank test; Figure 4C). Along the rostrocaudal axis, the mean value of full-width at half maximum for layer 5 was 1286.8 μm (SD = 131.9 μm), and 1473.7 μm (SD = 161.9 μm) for layer 6 (n = 10, p-value = 0.0020, Wilcoxon signed rank test; Figure 4D). Thus, it appears that layer 6 corticocollicular neurons occupy a significantly broader area in the cortex than layer 5, suggesting that these layer 6 neurons may route distinct forms of information from cortical areas surrounding the primary AC to the IC. In addition, the peak of layer 5 corticocollicular cells was found to be displaced by 99.9 μm dorsally and 425.25 μm caudally relative to layer 6 corticocollicular cells, (n = 10, p-value = 0.0020, Wilcoxon signed rank test; Figure 4E), which aligns the peak of the of the layer 5 corticocollicular distribution with the center of the lemniscal AC areas.

**Figure 4.**
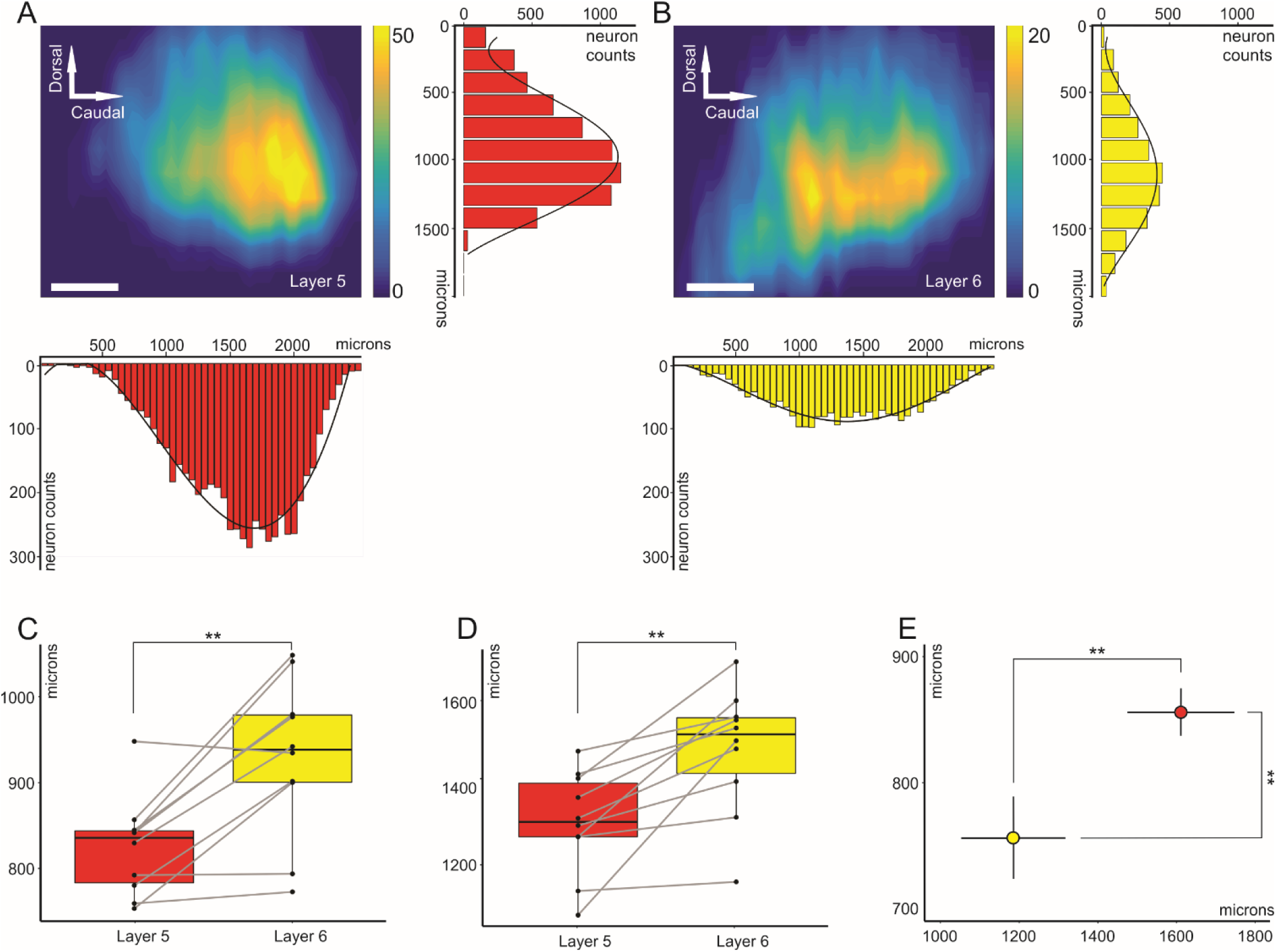
Partially segregated distributions of layer 5 and layer 6 corticocollicular neurons in mouse AC. (A) Distribution of layer 5 corticocollicular neurons in the neocortex, also showing the summation along the rows, which results a dorsoventral distribution, and summation along the columns – rostrocaudal distribution. (B) Distribution of layer 6 corticocollicular neurons from the same animal. (C) Comparison of full-width at half maximum measures for layers 5 and 6 in the dorsoventral direction and (D) rostrocaudal direction. (E) Estimated peaks of layer 5 (red) and layer 6 (yellow) corticocollicular distributions. The peak for layer 6 is shifted more ventrally and rostrally compared to layer 5. The error bars represent 95% confidence intervals. Scale bar = 250 μm.

Given the global anatomical differences between layer 5 and 6 corticocollicular neuronal distributions described above, we asked whether some of the more rostrally- and ventrally-located layer 6 corticocollicular cells were in acoustically-responsive zones. To answer this question, the left AC in GCaMP6s-Thy1 transgenic mice was mapped by imaging responses to amplitude-modulated pure tones at 5, 10, 20 and 30 kHz using wide-field transcranial optical imaging with blue light (Figure 5, panels A and B). Four auditory subfields could be identified reliably as described by previous investigators (Issa, Haeffele et al. 2014), with A1 and AAF organized tonotopically, converging in high frequency regions near weakly tonotopic secondary AC (A2). The ultrasonic field (UF) was located dorsally, and only responses to frequencies above 20 kHz were present in this field (Figure 5 B, C). After characterization of the sound-evoked responses, Fluorogold was injected into left IC of these mice, complete reconstructions of layer 5 and 6 corticocollicular distributions were obtained (Figure 5 D, E), and previously recorded responses were overlaid with layer 5 and 6 distributions (Figure 5 F, G).

**Figure 5.**
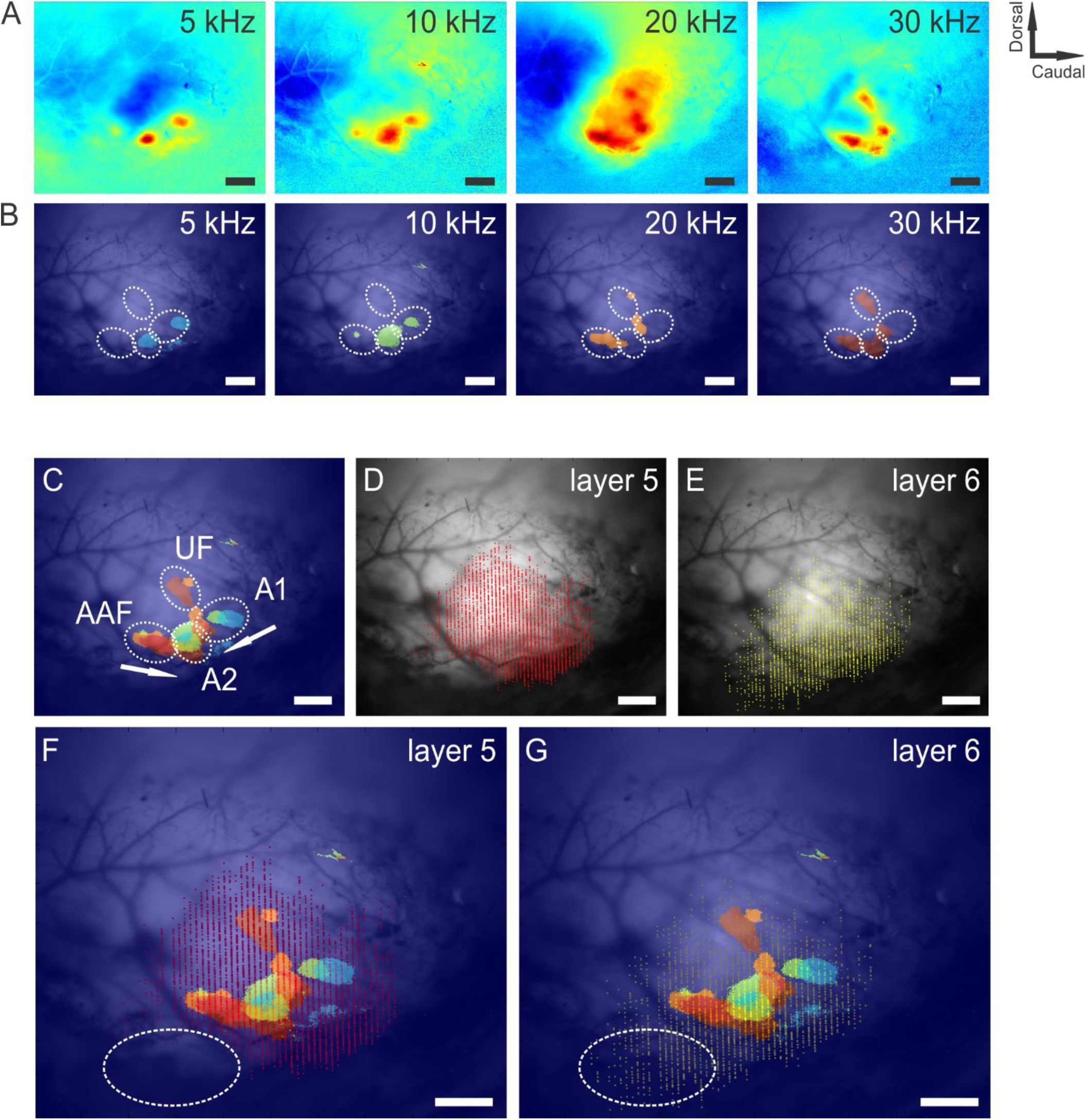
Functional maps and corresponding distributions of layer 5 and layer 6 corticocollicular neurons in mouse AC. *ΔF/F* responses to 100% amplitude-modulated pure tones in mouse AC (A). A threshold level at 2.5 standard deviations was set to responses in A to display the peaks of stimulus-evoked cortical activity (B). Combination of cortical responses at threshold level from (B) identifies different auditory regions: tonotopically organized A1 and AAF, as well as A2 and UF (C). Reconstructions of layer 5 (D) and layer 6 (E) corticocollicular neurons for the same mouse. (F) and (G) are overlays of the tonotopic map and corticocollicular reconstructions. Notice the absence of sound-evoked activity in the rostroventral to AC regions (highlighted in oval). **Scale bar** = 500 μm.

It was observed that the rostroventral cortical area containing primarily layer 6 corticocollicular neurons appeared outside the main acoustically-responsive regions, highlighted with an oval in Figures 5F and G. To determine whether this apparently acoustically-unresponsive zone would be responsive to more meaningful sounds, the same approach was applied to examine the responses of the AC to ethologically relevant sounds (Grimsley, Monaghan et al. 2011, Grimsley, Sheth et al. 2016). Several different species-specific mouse calls were used as stimuli. Again, after obtaining the reconstructions of layer 5 and 6 corticocollicular cells and overlaying this reconstruction with functional mapping, it was found that the rostroventral to the AC area enriched in layer 6 corticocollicular cells was outside sound-responsive functional cortical areas (Figure 6).

**Figure 6.**
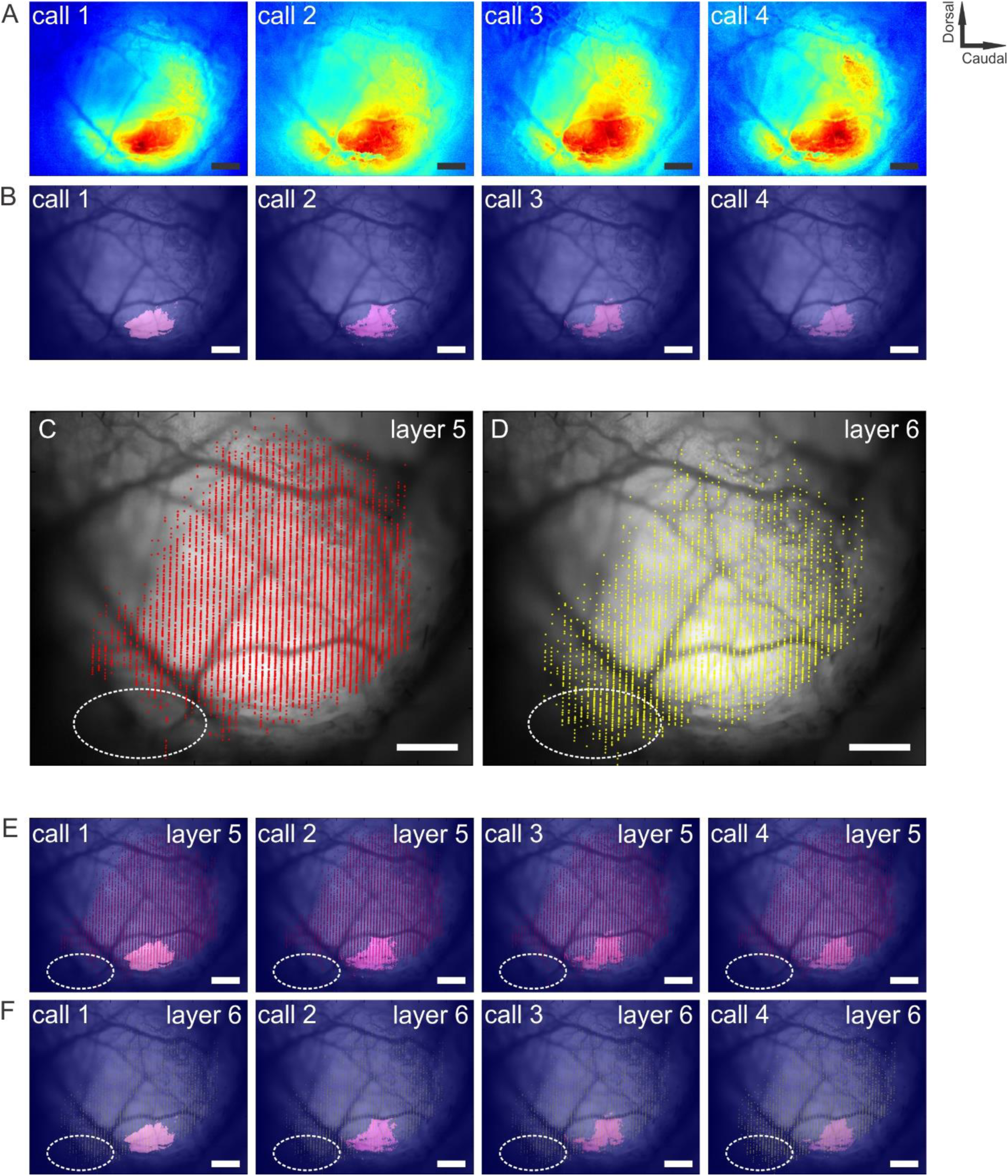
Functional responses to ethologically-relevant sounds and corresponding distributions of layer 5 and layer 6 corticocollicular neurons in mouse AC. *ΔF/F* responses to four classes of species-specific calls (A). Call 1 and 2 are low-frequency stress calls, call 3 is a restraint stress-induced medium-frequency call, and call 3 is a mating call. A threshold level at 2.5 standard deviations was set to responses in A to display the peaks of stimulus-evoked cortical activity (B). Distributions of layer 5 and 6 corticocollicular cells (C, D). Overlays of functional maps from (B) with layer 5 (E) and layer 6 (F) corticocollicular cells. **Scale bar** = 500 μm.

To further characterize this rostroventral region containing isolated layer 6 corticocollicular projections, Alexa-594-conjugated CTb was injected into this region to label its inputs. The distribution of the inputs from one of these experiments is presented in Figure 7. Consistent with the *in vivo* neuroimaging findings, the rostroventral region received the majority of its input from non-auditory structures, including the lateral nucleus of the amygdala (LA), nucleus of the brachium of the IC (NB), posterior limiting nucleus of the thalamus (POL), suprageniculate nucleus (SGN), and posterior complex of the thalamus (PO). Of note, there is essentially no input from the MGB, which is the main thalamic nucleus responsible for auditory processing. These connectivity findings are indicative of the non-auditory nature for this cortical area containing isolated layer 6 corticocollicular neurons.

**Figure 7.**
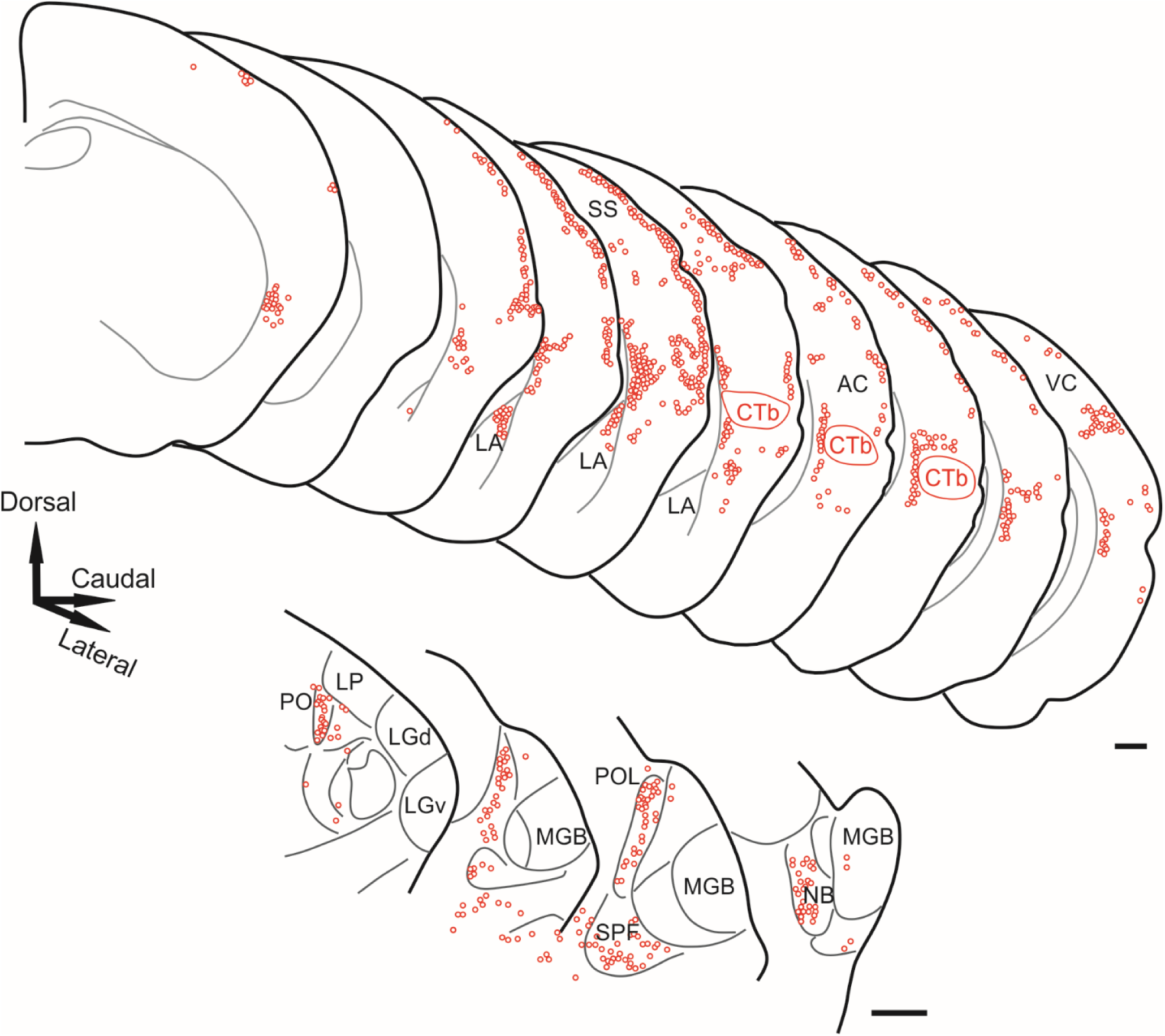
Inputs to the cortical area containing layer 6 but not layer 5 corticocollicular neurons. The top panel shows the injection site (CTb) and cortical areas where retrogradely labeled neurons were found. A significant portion of inputs came from the lateral nucleus of the amygdala (LA), as well as somatosensory and visual cortices (SS and VC, respectively). The bottom panel shows the distribution of neurons in the thalamus and associated structures. The majority of labeled neurons were found in the posterior complex (PO), posterior limiting nucleus (POL), subparafascicular nucleus (SPF) and nucleus of the brachium of the IC (NB), but not the medial geniculate body (MGB) Scale bar = 500 μm.

Layer 5 and layer 6 corticofugal neurons have been shown to have differential patterns of projections in other systems such as the corticothalamic pathway (Llano and Sherman 2008). These anatomical differences are also postulated to have functional significance in forward propagation of sensory information via the higher order thalamic nuclei (Llano and Sherman 2009, Theyel, Llano et al. 2010). Given the presence of functional neuroanatomical differences with respect to the cortical projections to thalamus, we next aimed to examine the termination patterns of layer 5 and 6 corticocollicular neurons in the mouse IC.

Rbp4-cre mouse line was successfully used by other researchers for labeling of layer 5 corticocollicular neurons (Xiong, Liang et al. 2015, Asokan, Williamson et al. 2018). Using this transgenic line and the following combination of AAVs, it was possible to achieve a significant separation between layer 5 and layer 6 corticocollicular termination patterns at the level of the IC. A mixture (500 nL) of equal concentrations of AAV9.CB7.CI.mCherry.WPRE.rBG to nonspecifically label corticocollicular projections with mCherry and AAV9.CAG.Flex.eGFP.WPRE.bGH to label Rbp4+ neurons with eGFP, were injected into the AC of two Rpb4-Cre mice. The mice were perfused two to three weeks later, and the viral expression was examined at levels of the AC and IC using confocal microscopy.

As expected, the nonspecific AAV construct was expressed throughout all layers of the AC, while the Cre-dependent AAV vector was only expressed in layer 5 (Figure 8). At the level of the IC, two kinds of labeled terminals could be identified: yellow terminals (eGFP + mCherry), which come from layer 5 neurons, and red terminals (mCherry) only, which arise from layer 6. Figure 8 displays a panel of coronal sections throughout the mouse IC with the patterns of expression. Consistent with numerous previous findings, the corticocollicular neurons primarily projected to the dorsal (DCIC) and lateral (LCIC) cortices of the IC. However, a layer-specific difference was found in the central nucleus of the IC (CNIC). This nucleus had a large number of yellow layer 5 terminals present, but not red-only layer 6 (two-sample Kolmogorov-Smirnov test, p-value = 5.032e-13) The overall distributions of size or number of layer 5 and 6 terminals did not vary significantly in the LCIC (two-sample Kolmogorov-Smirnov test, p-value = 0.29), or the DCIC (two-sample Kolmogorov-Smirnov test, p-value = 0.062). The layer 5 corticocollicular terminals in the CNIC also showed an axosomatic innervation pattern (Figure 9 B), which was supported by repeating this injection of AVV’s in a different Rpb4-positive mouse and immunostaining for neuronal marker NeuN (Figure 10).

**Figure 8.**
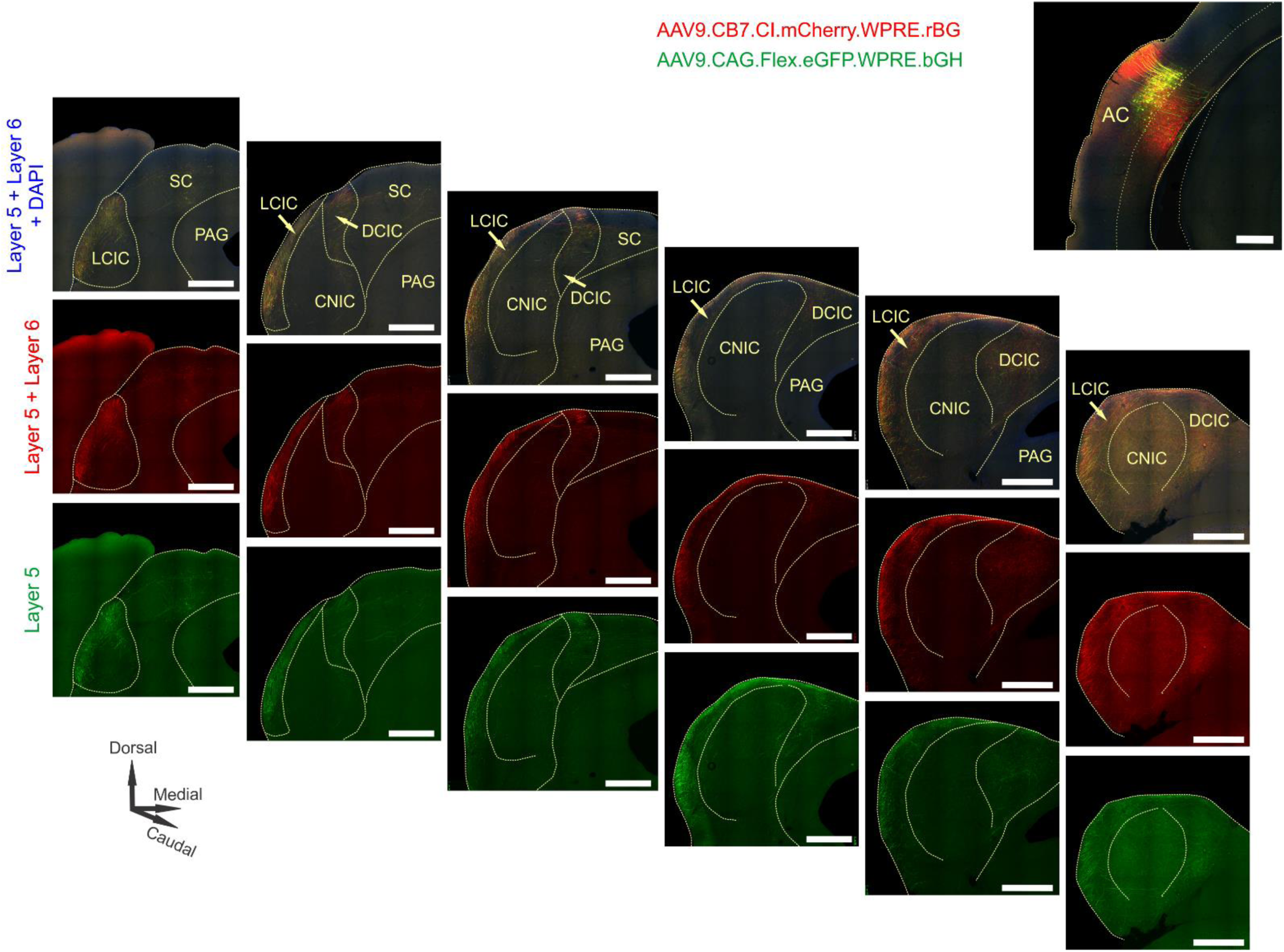
Corticocollicular termination patters at the level of the IC. Both layer 5 and layer 6 corticocollicular terminals from the AC were densely distributed in the LCIC and DCIC. The green panel (e.GFP expression) shows layer 5 terminals only. The middle panel (m.Cherry) is a combined labeling of layers 5 and 6. The top panel shows the terminals from both layers. The yellow color is the labeling from layer 5, while purely red terminals are dominated by layer 6. Top right - injection site in the AC, with the dashed yellow line showing the border between cortical layers 5 and 6. Scale bar = 500 μm.

**Figure 9.**
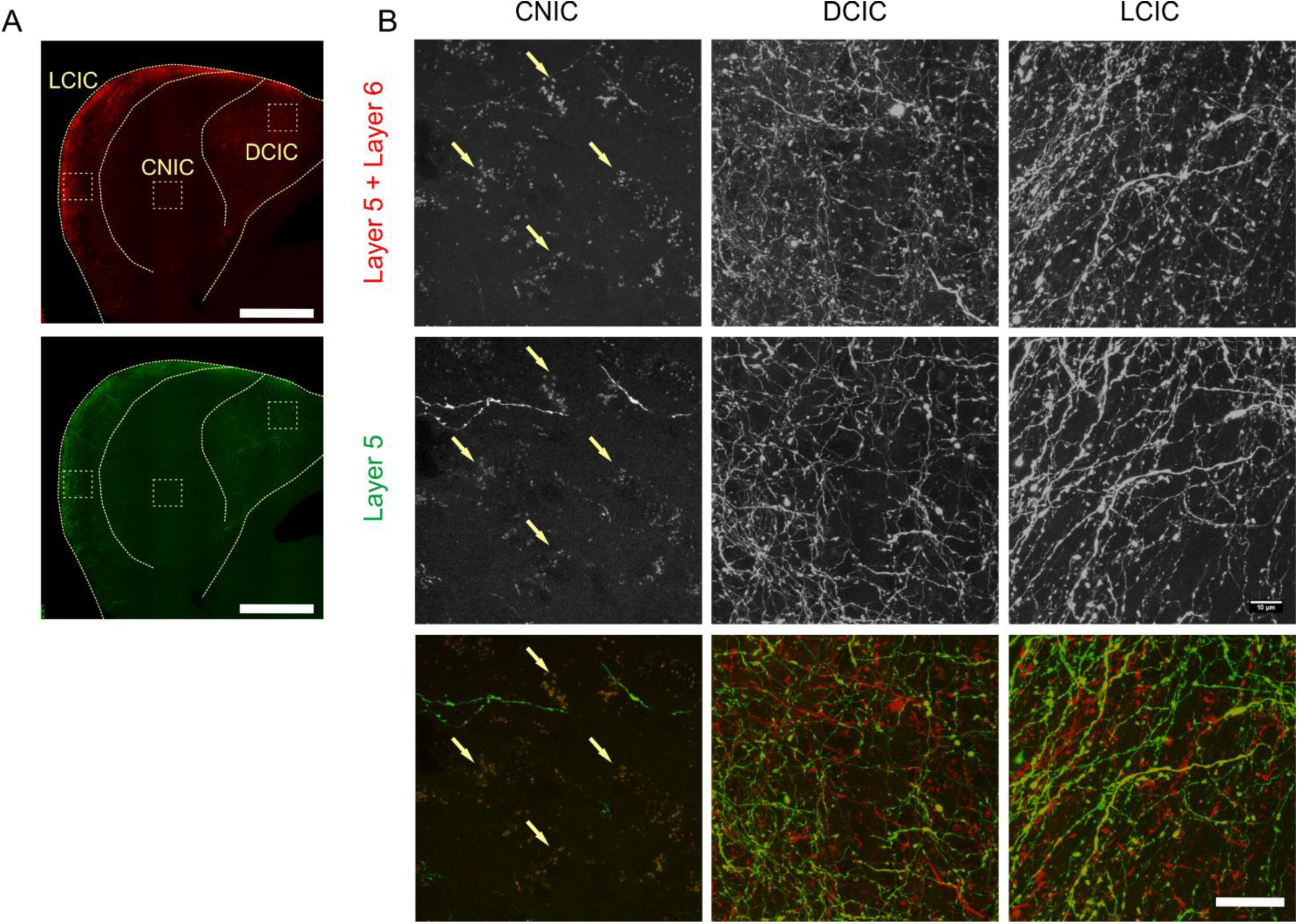
Details of the corticocollicular termination patterns in the subdivision of the IC. (A) AAV9 expression in layer 6 + layer 5 and in layer 5 in the Rbp4 mouse. The insets are regions whose higher resolution details are shown in B. (B). Although the CNIC received significantly less corticocollicular input overall, consistent with previous findings, a number synaptic boutons (arrows pointing) emanating from layer 5 were found in this subdivision, while layer 6 boutons were virtually absent. Scale bar = 500 μm in (A) and 20 μm in (B).

**Figure 10.**
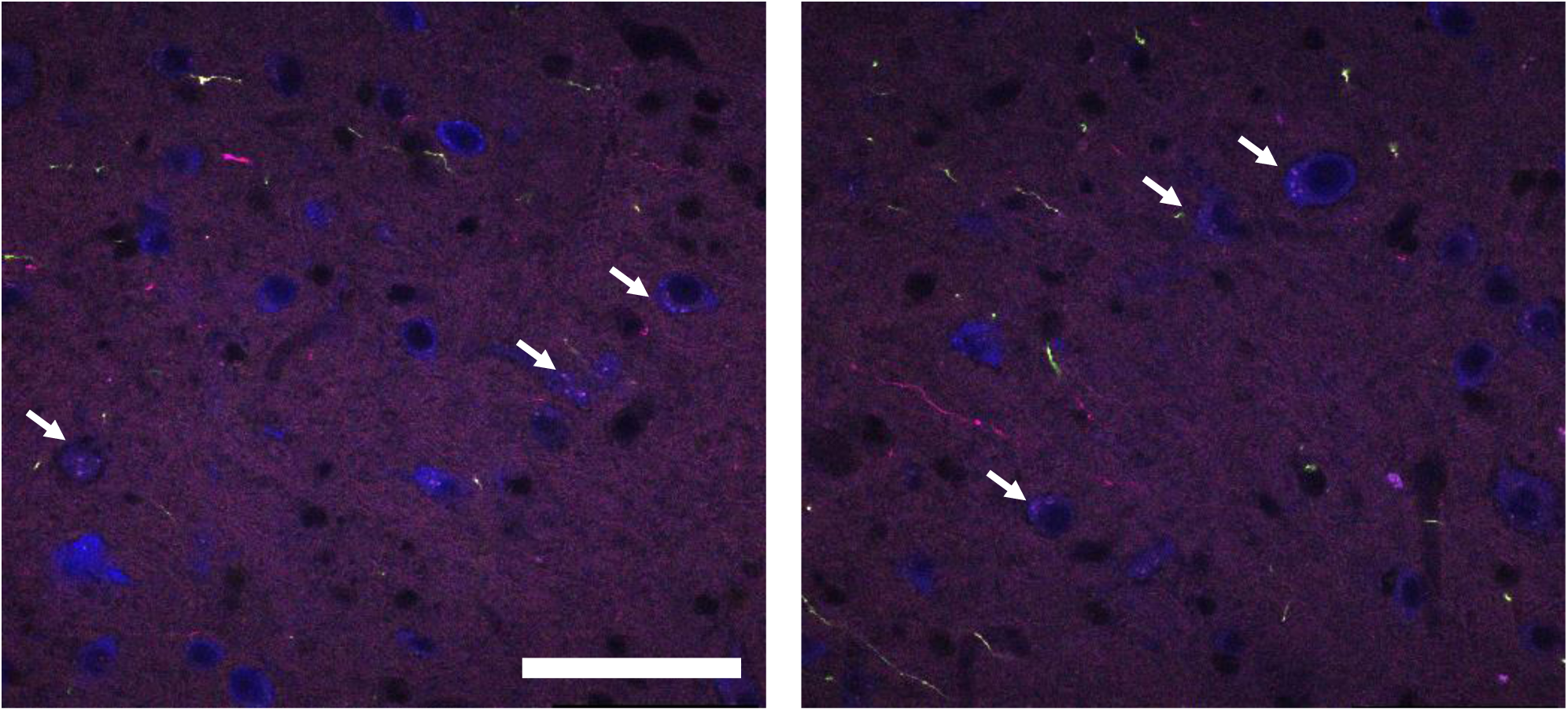
Layer 5 corticocollicular terminals contact the cell bodies of neurons in the CNIC. A single slice of a confocal stack showing AAV9 expression in layer 6 + layer 5 and in layer 5 in the Rbp4 mouse CNIC stained for NeuN. Layer 5 axosomatic synaptic terminals (arrows pointing) were found in this subdivision of the IC. Scale bar = 50 μm. Z-step = 500 nm.

To determine the specific distribution of layer 6 corticocollicular terminals, a Cre-OFF virus was injected into the primary AC of mice expressing Cre-recombinase in both Rbp4+ and CLGT+ cells in layer 5. Expression of the virus was seen in layer 6 of the AC (Figure 11A) and in the medial geniculate body (Figure 11B). In the IC, there was label seen in the most superficial portions of the rostral lateral cortex (Figures 11C and D).

**Figure 11:**
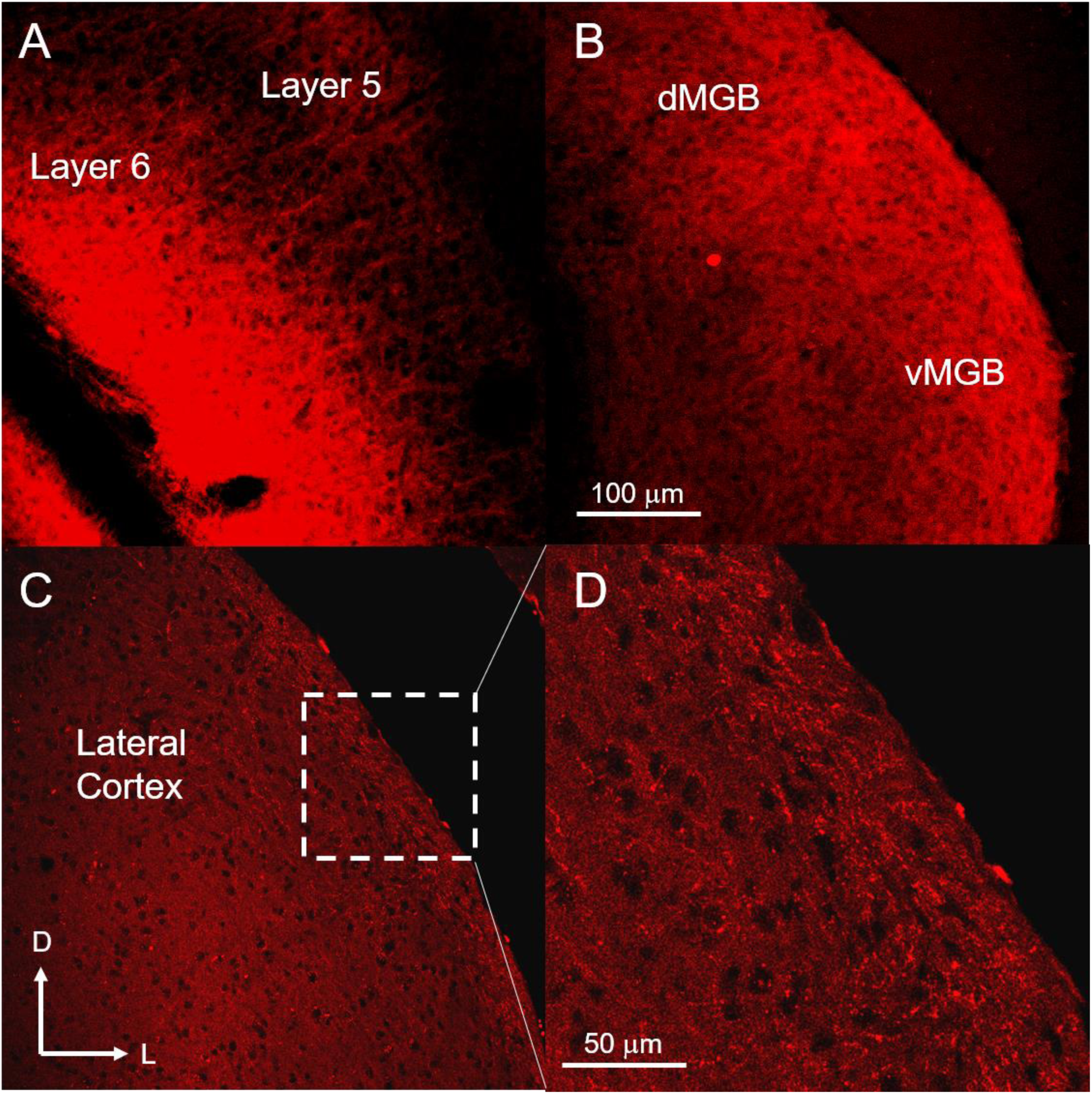
Layer 6 targets the surface of the lateral cortex of the IC. A) Injection site of the Cre-OFF virus into the auditory cortex. B) Label is seen in the dorsal and ventral medial geniculate body (dMGB, vMGB). C) Label is seen on the most superficial parts of the lateral cortex of the inferior colliculus. D) Expansion of the box in C highlighting the small terminals on the surface of the lateral cortex.

## Discussion

In this study, it was shown that the distributions of layer 5 and layer 6 corticocollicular neurons are partially but significantly non-overlapping with respect to each other and the cortical areas from which they originate. In particular, it was found that the IC receives heavy input from layer 6 from a cortical region rostral and ventral to the lemniscal AC, as indicated by staining for PV and SMI32. The peaks and widths varied for layer 5 and layer 6 corticocollicular neuronal distributions. Layer 6 corticocollicular neurons encompassed a broader area of the cortex than layer 5, and its peak shifted more rostrally and ventrally compared to layer 5. This organization implies that different cortical areas may modulate the IC either through layer 6 corticocollicular neurons or both layers 5 and 6.

Layer 5 and layer 6 corticocollicular neurons appear to be differentially distributed with respect to functionally mapped areas of the AC. More specifically, an area of the cerebral cortex located rostrally and ventrally to the mouse AC contains layer 6 but not layer 5 corticocollicular neurons. This cortical region does not respond to sound stimuli such as pure tones of different frequencies, neither is it responsive to more ethologically relevant sounds such as species-specific calls. It is possible that imaging the AC in awake animals may have elicited acoustically-driven responses from this region. However, the maps elicited in the current study are similar to the distributions of acoustically-responsive areas seen in previous studies in awake animals. In the current study, four regions of the mouse AC could be distinguished based on their responses to pure tones. A1 and AAF were tonotopically organized consistent with previous reports (Stiebler, Neulist et al. 1997, Tohmi, Takahashi et al. 2009, Issa, Haeffele et al. 2014). The ultrasonic frequencies pure tones activated neurons in the UF, while all pure tone stimuli also activated A2, located ventrally to A1 and AAF gradient convergence. These regions overlapped more so with the distributions of layer 5 corticocollicular neurons, and less with the corticocollicular neurons emanating from layer 6. Four types of mouse-specific calls used in this study activated neurons located primarily in A2. Again, this activation did not overlap with the cortical area containing only layer 6 but not layer 5 corticocollicular cells. Finally, this area did not receive input from any subdivisions of the MGB. The latter finding distinguishes this region from the insular auditory field, which received input from the ventral division of the MGB. Instead, intralaminar nuclei were the main source of thalamic input to this area. The lateral nucleus of the amygdala also sent a heavy projection to this area, as well as upper-layer neurons found in somatosensory and visual cortices. These findings suggest that only layer 6 corticocollicular neurons are found in brain regions that funnel a broad spectrum of visual, somatosensory and limbic information, potentially to modulate IC function in response to multisensory and emotional-stimuli.

Differences between layer 5 and 6 corticocollicular distributions found in this study and previous *in vitro* electrophysiological recordings from these neurons point to different plausible functions of these layers in auditory cortical and midbrain processing (Slater, Willis et al. 2013). Consistent with previous findings (Herbert, Aschoff et al. 1991, Budinger, Heil et al. 2000, Bajo, Nodal et al. 2007), in our study layer 5 corticocollicular neurons also appeared to be confined to tonotopically-organized lemniscal auditory fields (A1 and AAF) as confirmed by PV and SMI-32 immunostaining and imaging. Thus, these corticocollicular neurons may be the main drivers of frequency shifts and changes in tuning duration observed in the IC upon AC stimulation. Unlike layer 5, the non-overlapping area of layer 6 corticocollicular neurons found rostrally and ventrally may be important for integrating information from other sensory modalities. Likely candidates for the origin of these projections are the perirhinal area and temporal association area (Suzuki 1996). These layer 6 corticocollicular neurons would be preferentially positioned to receive integrated multisensory information from upper layers of these association areas and then route these signals to the IC, thus adjusting neural activity in the auditory midbrain according to animal’s behavioral needs. Further anatomical, physiological and behavioral experiments are needed to address layer-specific differences in the corticocollicular system and their roles in auditory information processing.

## Acknowledgments

The authors thank Dr. Jasmine Grimsley for providing recordings of mouse calls and assistance in their use. This work was supported by DC013073 to D.A.L.

